# The IRE1a-XBP1 pathway promotes T helper cell differentiation by resolving secretory stress and accelerating proliferation

**DOI:** 10.1101/235010

**Authors:** Jhuma Pramanik, Xi Chen, Gozde Kar, Tomás Gomes, Johan Henriksson, Zhichao Miao, Kedar Natarajan, Andrew N. J. McKenzie, Bidesh Mahata, Sarah A. Teichmann

**Affiliations:** Wellcome Trust Sanger Institute, Wellcome Genome Campus, Hinxton, Cambridge, CB10 1SA, United Kingdom; EMBL-European Bioinformatics Institute, Wellcome Genome Campus, Hinxton, Cambridge, CB10 1SD, United Kingdom; MRC Laboratory of Molecular Biology, Cambridge Biomedical Campus, Francis Crick Avenue, Cambridge CB2 OQH, United Kingdom; Theory of Condensed Matter, Cavendish Laboratory, 19 JJ Thomson Ave, Cambridge CB3 0HE, United Kingdom.

**Keywords:** Th2 lymphocyte, XBP1, Genome wide XBP1 occupancy, Th2 lymphocyte proliferation, ChIP-seq, RNA-seq, Th2 transcriptome

## Abstract

The IRE1a-XBP1 pathway, a conserved adaptive mediator of the unfolded protein response, is indispensable for the development of secretory cells. It maintains endoplasmic reticulum homeostasis by facilitating protein folding and enhancing secretory capacity of the cells. Its role in immune cells is emerging. It is involved in dendritic cell, plasma cell and eosinophil development and differentiation. Using genome-wide approaches, integrating ChIPmentation and mRNA-sequencing data, we have elucidated the regulatory circuitry governed by the IRE1a-XBP1 pathway in type-2 T helper cells (Th2). We show that the XBP1 transcription factor is activated by splicing *in vivo* in T helper cell lineages. We report a comprehensive repertoire of XBP1 target genes in Th2 lymphocytes. We found that the pathway is conserved across cell types in terms of resolving secretory stress, and has T helper cell-specific functions in controlling activation-dependent Th2 cell proliferation and regulating cytokine expression in addition to secretion. These results provide a detailed picture of the regulatory map governed by the XBP1 transcription factor during Th2 lymphocyte activation.

## Introduction

T helper (Th) cells (CD4^+^ T cells) are central to the adaptive immune response, immune-tolerance and potentiate innate immune response pathways (Walker and McKenzie, 2017; Zhu et al., 2010). Therefore, these cells are key players in infections, allergies, auto-immunity and anti-tumor immune responses. Depending upon the immunogen or allergen (*e.g.* infection, commensal microorganism, or self-antigen), naive T helper cells become activated, proliferate and are able to differentiate into several subtypes, such as Th1, Th2, Th17, and regulatory T cell (Treg). This Th subtype classification is based on their differential expression of cytokines and key lineage specific transcription factors (Murphy and Reiner, 2002; Zhu et al., 2010). Th2 lymphocytes secrete their characteristic cytokines IL4, IL5, IL10 and IL13. These secretory cells are involved in worm parasite expulsion, exaggerate allergies and asthma, potentiate pregnancy (Wegmann et al., 1993) and suppress anti-tumor immunity (Ellyard et al., 2007). Transcription factors that are involved in differential production and regulation of cytokine genes, for example GATA3 in Th2, are well studied. However cytokine gene expression is only one aspect of the T helper cell differentiation process. The ability to rapidly proliferate is another key attributes of T helper lymphocytes (Figure 1A), and the full regulatory circuitry controlling these processes is still incompletely understood.

**Figure 1:**
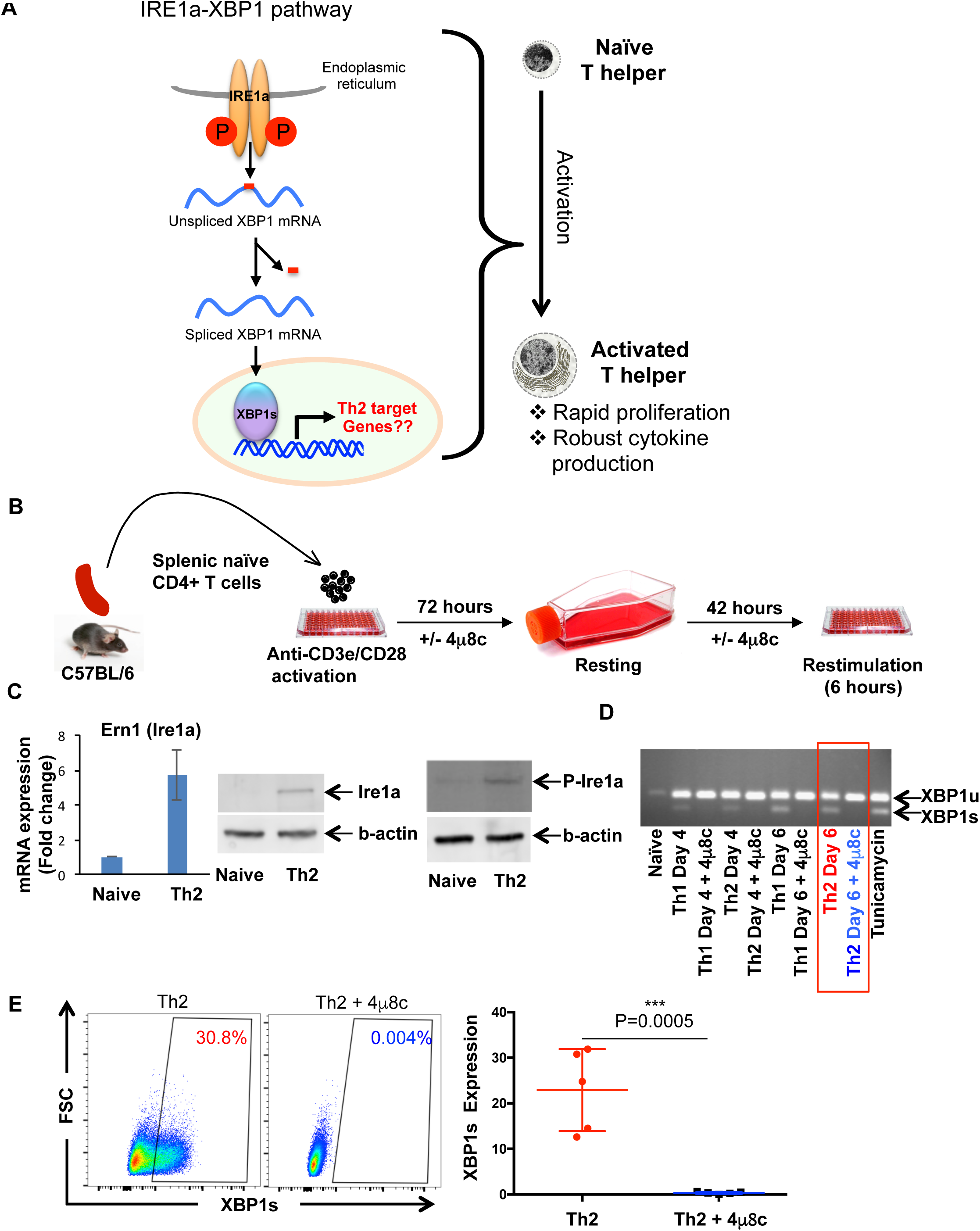
*T helper cells up-regulate the IRE1a-XBP1 pathway during activation.* A. Schematic representation of the hypothesis. In this study we are asking what role does the IRE1a-XBP1 pathway play during T helper cell activation. T helper cell activation is a dramatic transformation from a quiescent cell state to a rapidly proliferative and highly protein productive/secretive cellular state. B. Overview of the experiment. Splenic naïve T cells were purified by negative selection and activated in anti-CD3e/C28 antibody coated plates under Th1 or Th2 conditions for 72 hours, rested for 42 hours and restimulated on anti-CD3e/CD28 antibody coated plate. C. Naïve T helper cells and *in vitro* differentiated Th2 lymphocytes were analyzed for IRE1a mRNA expression by qRT-PCR (left panel), protein expression by western blot (middle panel), and phosphorylated IRE1a (P-IRE1a) by western blot (right panel). D. Naïve T cells were cultured under Th1 or Th2 differentiation conditions in the presence or absence of IRE1a inhibitor (4μ8c. *In vitro* activated T helper cells were analyzed by RT-PCR using a pair of primers that discriminate the cDNA derived from spliced and unspliced form of XBP1 mRNA. E. *In vitro* differentiated Th2 cells were stained with fluorescent dye conjugated anti-XBP1s specific antibody and analyzed by flow cytometry. One representative FACS profile is displayed (left panel) and the graph containing all results (n=5) is shown in “right panel”.

Proliferation is required for clonal expansion, which forms the basis of the adaptive immune response (Bird et al., 1998; Gett and Hodgkin, 1998). The Gata3/RuvB-like protein 2 (Ruvbl2) complex was shown to be a key regulatory driver of Th2 cell proliferation (Hosokawa et al., 2013), and several other transcription factors, such as Stat6, are implicated in the regulatory circuitry controlling T helper cell proliferation and differentiation. Additional transcription factors are likely to be involved in regulating this highly organized, complex process.

At the cell biological level, to synthesize, fold and secrete proteins, including cytokines, activated T helper cells need to contain a well-differentiated endoplasmic reticulum (ER) and protein secretory machinery. It is an open question how activated T helper cells meet this protein folding and secretory demand. Secretory cells (e.g. pancreatic β-cell, acinar cells) address this challenge by up-regulating the unfolded protein response (UPR) pathway triggered by the accumulation of unfolded proteins in the endoplasmic reticulum (ER) (Calfon et al., 2002; Frakes and Dillin, 2017; Hetz, 2012). Three ER membrane-resident sensors, the endonuclease IRE1a (encoded by *ERN1* gene), the kinase PERK and the cleavable precursor of the transcription factor ATF6 coordinate the process. Among these three, the IRE1a-XBP1 pathway is the most evolutionary conserved pathway (Figure 1A) (Hetz and Papa, 2017; Kaser and Blumberg, 2010). During ER stress, the kinase, IRE1a, oligomerizes, autophosphorylates, and uses its endoribonuclease activity to splice a 26-nucleotide fragment from the unspliced XBP1 mRNA (XBP1u). This then results in the functional spliced form of the transcription factor XBP1 (XBP1s) (Hetz et al., 2011). XBP1s regulates the expression of numerous target genes involved in ER biogenesis. Its role has been studied in secretory cells, such as pancreatic acinar cells, plasma cells and dendritic cells (DC). In these cell types XBP1 occupies chromatin and controls gene expression in a cell-type specific manner (Acosta-Alvear et al., 2007). This suggests that XBP1 may play a role in diverse cell types. Therefore we set out to investigate its specific function in CD4^+^ T lymphocytes (Figure 1A).

The role of the IRE1a-XBP1 pathway in immunity and inflammation is now emerging (Bettigole and Glimcher, 2015; Brucklacher-Waldert et al., 2017; Grootjans et al., 2016; Hotamisligil, 2010; Janssens et al., 2014). The pathway has been described in dendritic cells, plasma cells, CD8^+^ T cells and eosinophil development and differentiation (Bettigole et al., 2015; Brunsing et al., 2008; Iwakoshi et al., 2007; Osorio et al., 2014; Thaxton et al., 2017; Todd et al., 2009). Interestingly, it has been reported recently, that the pathway causes cancer-associated immune suppression by causing dendritic cell dysfunction (Cubillos-Ruiz et al., 2015). The pathway is also involved in alternative activation of macrophages, and in obesity (Shan et al., 2017). Together, these reports suggest that the XBP1 transcription factor can contribute to a wide range of biological processes. IRE1a inhibitors (e.g. 4μ8c have been proposed as a treatment of cancer, by reinstating cancer immunity and eosinophilia by inhibiting eosinophil differentiation (Bettigole et al., 2015; Cubillos-Ruiz et al., 2017; Cubillos-Ruiz et al., 2015; Hetz et al., 2013). Here we test the role of the XBP1 transcription factor in regulating T helper cell activation through inhibition of the IRE1a-XBP1 pathway by the small molecule inhibitor 4μ8c

Using genome-wide approaches, integrating transcriptomic and XBP1 chromatin occupancy data, we elucidate the regulatory circuitry governed by the IRE1a-XBP1 pathway in Th2 lymphocytes. We found that the pathway observed in other cells is conserved in T helper cells in terms of secretory stress adaptation. Further, we show that XBP1 regulates genes that control diverse facets of Th2 cell physiology. In addition to resolving protein folding and secretory stress, it accelerates cell proliferation, and controls cytokine synthesis and secretion.

Our data provide a rich resource for investigating XBP1-regulated genes with genome-wide chromatin occupancy and expression, with a browsable online database at: http://data.teichlab.org

## Results and Discussion

### *T helper cells switch on the IRE1a-XBP1 pathway during* in vitro *activation*

Activated and differentiated T helper cells secrete an abundance of cytokines. Therefore, a well-developed secretory machinery is a prerequisite for cells to adapt to this secretory stress. RNA-seq data analysis predicts that when naïve T helper cells are activated and differentiated into Th2 cells, they upregulate expression of genes involved in the ER stress pathway (Supplementary Figure S1). Several factors that have previously been characterized as controllers of protein folding and secretion, including XBP1 itself, are upregulated during T helper cell differentiation.

To validate this prediction, and specifically investigate the involvement of the IRE1a-XBP1 pathway, we measured IRE1a mRNA and protein expression in T helper (Th) cells differentiated *in vitro* (Figure 1B). We found that both mRNA and protein level were upregulated in activated T helper cells (Figure 1C, left and middle panel). It is known that phosphorylation of IRE1a denotes its functional state. We observed that the protein is phosphorylated upon activation (Figure 1C, right panel).

Activated IRE1a splices the unspliced XBP1 (XBP1u) mRNA and produces a spliced XBP1 (XBP1s) mRNA isoform. We observed increases in the spliced form of XBP1 (XBP1s), both at mRNA and protein levels, upon T helper cell activation (Figure 1D, E). Specific inhibition of the IRE1a endonuclease activity by treating the cells with 4μ8c (Cross et al., 2012) abolished both the XBP1s mRNA and protein isoforms, confirming that the formation of the spliced form was dependent on IRE1a activity (Figure 1D, E).

These results confirm that the IRE1a-XBP1 pathway is conserved in T helper cells, and upregulated during T helper cell activation. Next, we set out to investigate whether this also holds *in vivo*.

### In vivo *activated T helper cells upregulate the IRE1a-XBP1*

To test whether the IRE1a-XBP1 pathway is operational in CD4^+^ T cells *in vivo,* we infected C57BL/6 mice with the helminth parasite *Nippostrongylus brasiliensis,* a well-established model of Th2-driven immune responses (Camberis et al., 2003; Mahata et al., 2014; Neill et al., 2010). After 7 days post-infection, we analyzed XBP1s protein expression in T helper cells by flow cytometry. We found T helper cells from worm-infected mice express significantly more XBP1s compared to uninfected control mice, suggesting an upregulation of the pathway (Figure 2A). We analyzed our previously published full-length single-cell RNA-sequencing data obtained from *N. brasiliensis* infections (Proserpio et al., 2016) as described in the "methods" section. We found T helper cells of immune challenged mice express XBP1s (Figure 2B).

**Figure 2:**
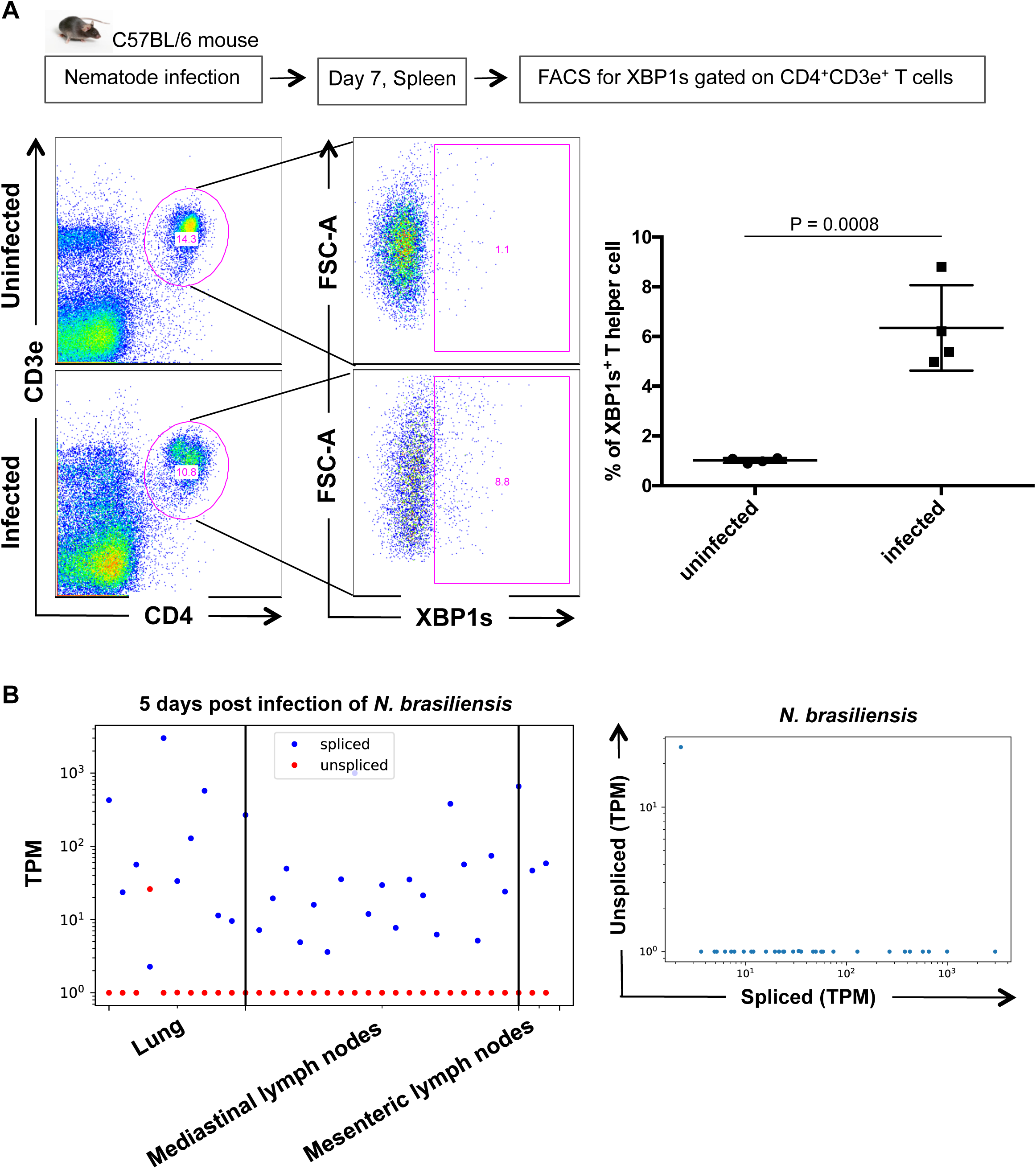
*T helper cells up regulate IRE1a-XBP1 pathway* in vivo *during infection.* A. Splenocytes from nematode (*Nippostrongylus brasiliensis)* infected mouse (7 days post infected) were stained with a PE conjugated anti-XBP1s antibody and analyzed by flow cytometry (Gating strategy: Singlet > live cells > CD4^+^CD3e+ > XBP1s+). One representative FACS profile is displayed (left panel) and the graph containing all results (n=4) is shown in the “right panel”. B. Our previously published single cell transcriptomic data obtained from mouse model of infection (i.e. *N. brasiliensis*) were reanalyzed to measure expression level (Transcript per million, TPM) of unspliced (red dots) and spliced (blue dots) form of XBP1 mRNA (left panel). Expression ratio of these two forms of XBP1 mRNA (right panel).

These results confirm that the pathway is active *in vivo.* Therefore, we set out to dissect the pathway using genome-wide approaches in Th2 lymphocytes.

### Genome-wide transcriptomic analysis of differential gene expression reveals IRE1a-XBP1 regulated genes

To capture a global gene regulatory role of the IRE1a-XBP1 pathway, we compared *in vitro* activated Th2 cells to cells with inhibited IRE1a endonuclease activity by adding 4μ8c into the cell culture media. We then compared the transcriptomes of activated Th2 lymphocytes with or without inhibition of the IRE1a-XBP1 pathway. Transcriptomes of 4μ8c treated and untreated Th2 cells were obtained by mRNA sequencing (RNA-seq). Quality control of the RNA sequencing data is shown in Supplementary Figure S2. Comparing transcriptomes of naïve and activated Th2 lymphocytes, we found that 10995 genes were differentially regulated upon Th2 activation. Inhibition of the IRE1a-XBP1 pathway by 4μ8c treatment resulted in differential expression of 3144 genes as compared to the untreated Th2 control (Figure 3A, Supplementary Figure S2 right panel). 2670 of these genes were involved in Th2 differentiation (Figure 3A). The pattern of differential expression reveals that these 3144 genes include 1693 up-regulated and 1451 down-regulated genes upon Th2 activation were found down regulated or up regulated respectively upon IRE1a inhibition (data not shown, but the similar pattern for protein folding and ER-stress related genes can be found at Figure 3B). Hierarchical clustering of the genes reveals the groups of genes up and down regulated upon 4μ8c treatment (Supplementary Figure S2, right). Detailed examination of these genes revealed many to be associated with the unfolded protein response and ER-stress, indicating a major impact of the IRE1a-XBP1 pathway (Figure 3B) on these biological processes. The complete list of differentially expressed genes can be found in Supplementary Table S1. Gene Ontology (GO) analysis of these differentially expressed genes upon 4μ8c treatment to Th2 cells (i.e. IRE1a-XBP1 pathway regulated genes) showed that they are enriched in the following biological processes: “Response to ER stress” (GO:0006950), “Regulation of signal transduction” (GO:0009966), “Cytokine production” (GO:0001816), “cell proliferation” (GO:0008283), “cell cycle” (GO:0007049) and Immune response (GO:0006955) (Figure 3C). These changes in the gene expression patterns upon IRE1a inhibition suggest extensive involvement of XBP1 transcription factor in Th2 activation and proliferation, as well as differentiation. Therefore, we set out to find the genome-wide chromatin occupancy patterns of the XBP1 transcription factor.

**Figure 3:**
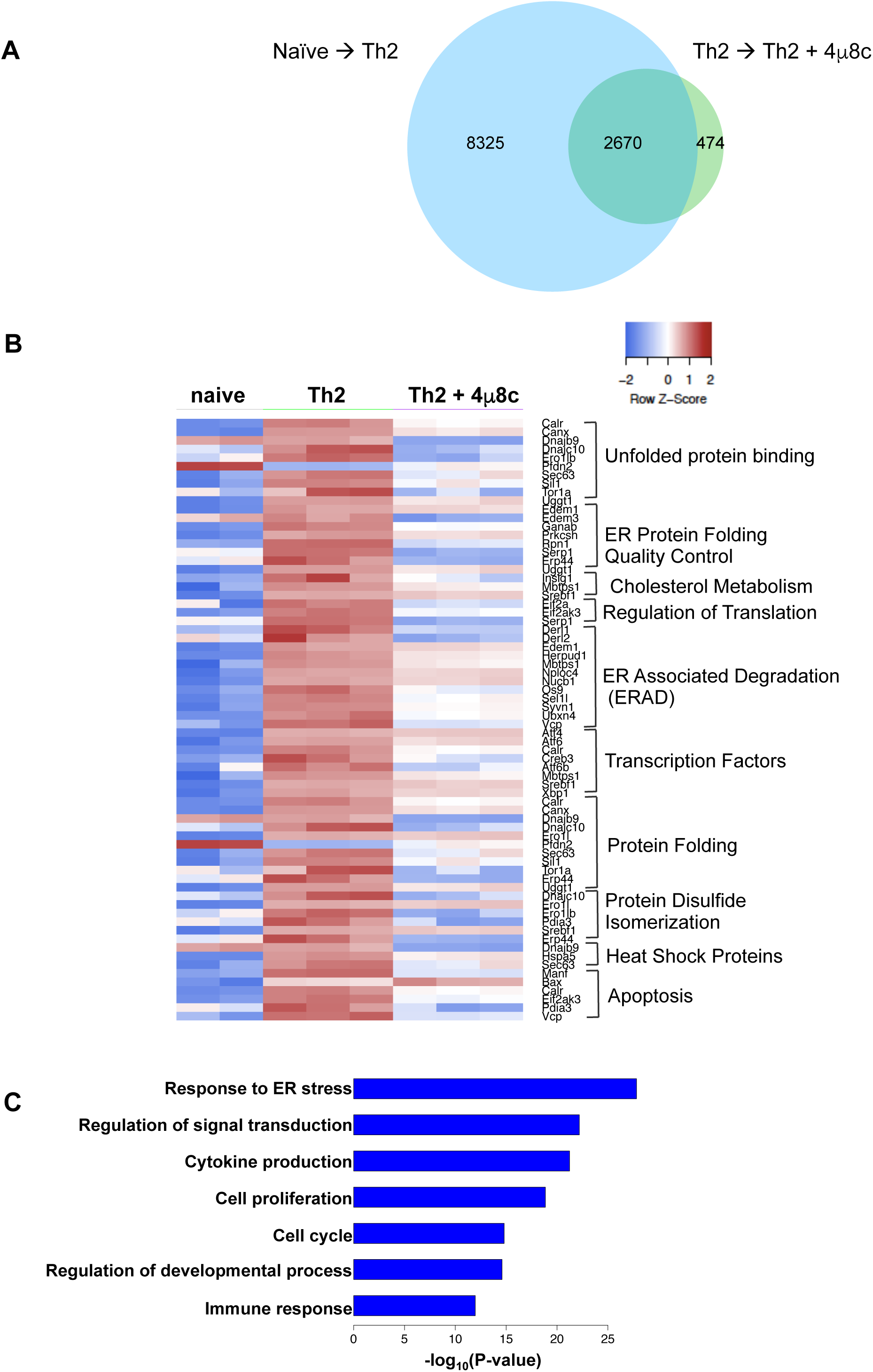
*Differential gene expression in Th2 due to the inhibition of IRE1a-XBP1 by* 4μ8c. Naïve T helper cells were activated under Th2 differentiation conditions in presence or absence of 4μ8c. Cells were activated in anti-CD3e and anti-CD28 antibody coated plates for 3 days, rested for 2 days, and reactivated in coated plates for 6 hours. The RNAseq data were analyzed for differential gene expression. A. Venn diagram showing the numbers of differentially expressed genes in different experimental conditions. “Naïve-> Th2” indicates the differentially expressed genes between naïve T helpers and Th2 cells. “Th2-> Th2+4μ8c” indicates the differentially expressed genes between untreated and 4μ8c treated Th2. A. Heatmap showing differentially expressed genes that are well known to be involved in resolution of ER stress imposed by unfolded protein response. B. Gene ontology (GO) analysis of the differentially expressed genes between Th2 and 4μ8c treated Th2.

### XBP1 ChIPmentation reveals XBP1 direct target genes in Th2 cells

To identify the genome-wide chromatin occupancy of XBP1, we performed ChIPmentation, a recently developed method that has been shown to be faster, more sensitive, and robust than traditional ChIP-seq approaches (Schmidl et al., 2015a), using a ChIP-grade antibody against XBP1. Two biological replicates were performed, and a total of 78 million reads were generated and mapped to the mouse genome. We identified 1,281 XBP1 binding peaks using MACS2 (Zhang et al., 2008) with a q-value less than 0.01.

As expected, binding peaks were identified around promoter regions in known XBP1 target genes, such as *Hspa5*, that encodes ER-chaperone protein BiP also known as Grp78, a master regulator of the unfolded protein response (UPR) (Figure 4A) (Lee et al., 2003). Interestingly, a binding event was also observed around the promoter of *Xbp1* itself (Figure 4A), indicating potential auto-regulation of Xbp1. The majority of the XBP1 binding peaks were located within promoter (36%) and intronic (35%) regions, and distal intergenic binding events (25%) were also frequently observed (Figure 4B). The genomic distribution of XBP1 peaks indicates that it binds both promoters and potential enhancers.

**Figure 4:**
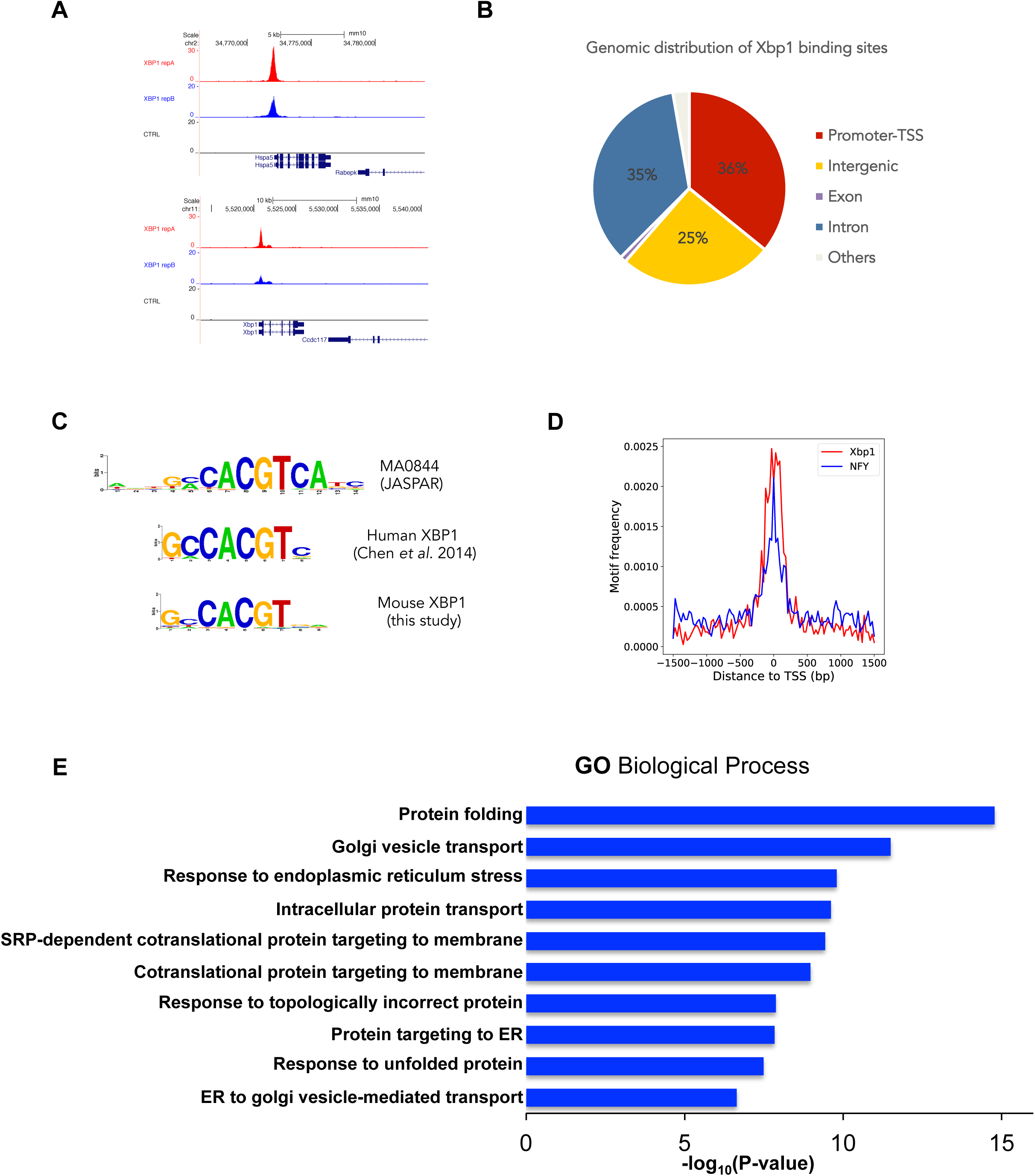
*Genome-wide chromatin occupancy of XBP1 transcription factor in Th2 lymphocyte.* XBP1 ChIPmentation was performed in *in vitro* differentiated Th2 cells to obtain genomewide XBP1 chromatin occupancy. A. Snapshot of XBP1 binding peaks around indicated representative genes from the UCSC genome browser. B. Genomic distribution of XBP1 binding peaks. The sector corresponding to the promoter includes sequences up to 1 kb upstream and 100 bp downstream from the TSS. C. Comparing the XBP1 motifs from the JASPAR database (top), ChIP-seq of the human breast cancer cell lines (middle), and mouse Th2 lymphocytes (bottom). D. Motifs frequencies of XPB1 and NF-Y around the binding peaks of XBP1. E. Top 10 biological processes GO terms enriched within XBP1 binding peaks analyzed by GREAT.

To further characterize the XBP1 regulome, we performed *de novo* motif discovery using HOMER (Heinz et al., 2010) to identify enriched DNA motifs within XBP1 binding regions. The top motif identified is the consensus sequence GCCACGT, which is almost identical to the human XBP1 binding motif defined in breast cancer cell lines (Figure 4C) (Chen et al., 2014). This indicates highly conserved binding specificities of XBP1 between human and mouse, and across cell types. The top motif enriched in our mouse data also resembles the XBP1 motif from the JASPAR database (Mathelier et al., 2016), again supporting the high quality of our ChIPmentation data. The second most enriched motif is the NF-Y binding motif (Supplementary Figure S3). Interestingly, the NF-Y motif has been frequently found around promoter regions of cell cycle genes, especially genes involved in G2/M cell cycle regulation (Chen et al., 2013; Muller et al., 2012). Both the XBP1 motif and the NF-Y motif co-occur around a subset of 65 XBP1 binding peaks (Figure 4D), indicating potential cooperation between XBP1 and NF-Y transcription factors to regulate a subset of target genes. The list of target genes that are potentially coregulated by XBP1 and NF-Y is displayed in Supplementary Table S2, and a complete list of targets is provided in Supplementary Table S3. This fact together with the observation from differentially expressed cell cycle/cell proliferation related genes (Figure 3C) prompted us to check the activation-induced proliferation of T helper cells (see later section below). The top five enriched motifs are displayed in Supplementary Figure S3. To investigate the functions of XBP1-bound genes, we used GREAT (McLean et al., 2010) to characterize XBP1 binding peaks. Most of the significant GO terms are related to protein folding and ER-stress (Figure 4E), which is consistent with the known biological role of XBP1.

Altogether the ChIPmentation experiments suggest a role of XBP1 in enhancing protein folding and secretion, as well as the proliferative capacity of Th2 cells.

### Integration of transcriptomic data and ChIP-seq data to unravel the XBP1-controlled gene regulatory network

To reveal the XBP1-regulated direct target genes and its transcriptional regulatory network, we integrated the genome-wide transcriptomic data and ChIPmentation data. A direct target gene is defined by its differential expression upon IRE1a inhibition (i.e. 4μ8c treatment) and XBP1 transcription factor occupancy at the gene locus. We found 435 direct target genes in Th2, of which 91 targets were previously reported as XBP1 direct target in other cell types (*i.e.* muscle, pancreatic β-cell and plasma cell) (Figure 5A). In this context, 344 genes can be considered as Th2-specific. XBP1 action over its direct targets has no defined direction, containing genes up and downregulated. The top 38 genes following either of these patterns are shown in Figure 5B, and the complete list can be found in Supplementary Table S4. The most significant identified GO terms are related to protein folding, ER-stress and secretory organelle homeostasis (Supplementary Figure S4A), which are consistent with its known biological roles, and also include novel Th2-specific targets.

**Figure 5:**
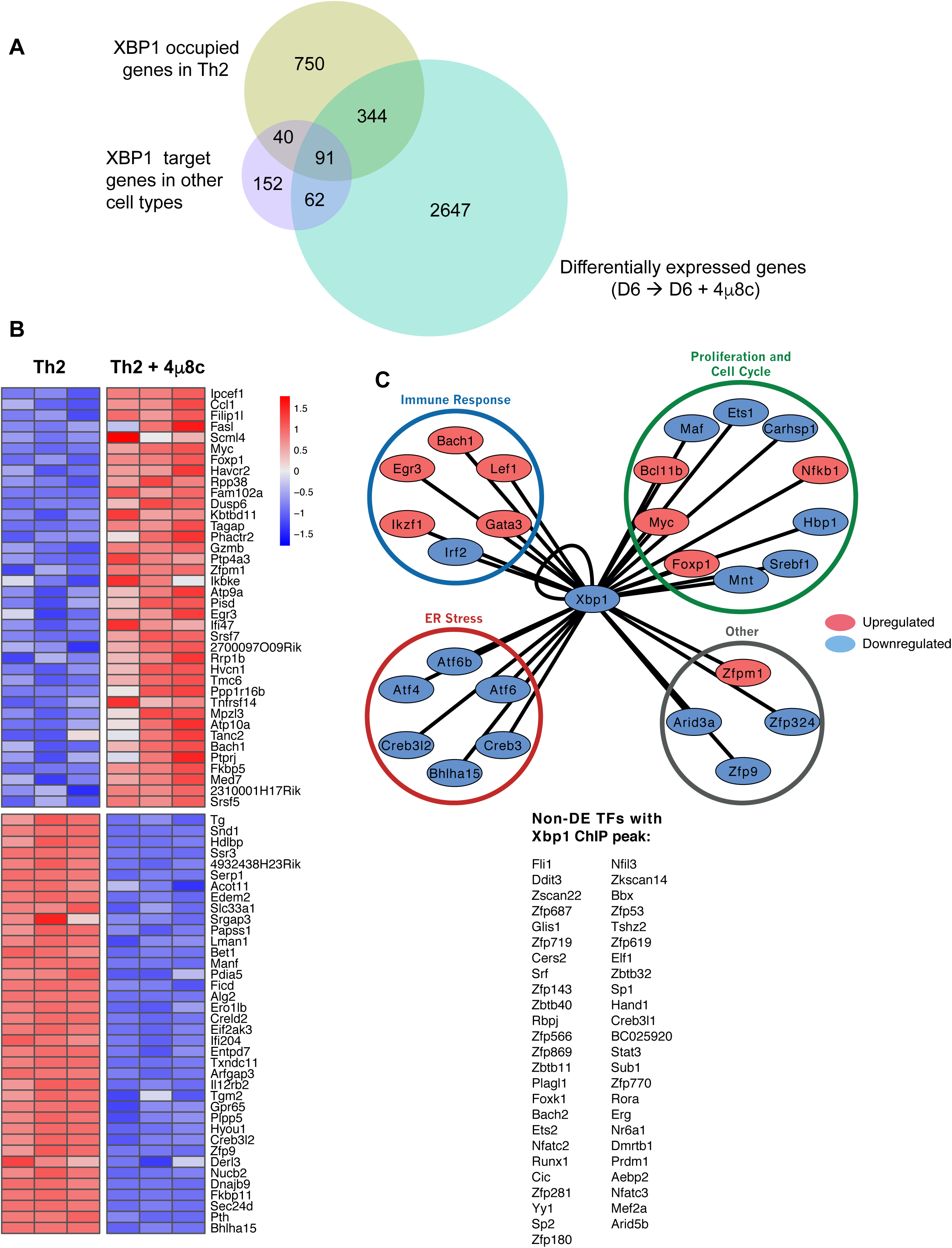
*Integration of ChIPmentation and RNA-seq data reveals XBP1 direct target genes and regulatory network.* A. Venn diagram comparing previously reported XBP1 target genes of other secretory cell types with the Th2 direct target genes of this study. XBP1 direct target genes of this study are those that are common in both “XBP1 occupied genes in Th2” and “Differentially expressed genes (Th2->Th2+4μ8c)” categories. The XBP1 direct target genes of B cell/plasma cell, skeletal muscle cells, pancreatic b-cells, were as observed by (Acosta-Alvear et al., 2007) and have been used here for comparison. B. Heatmap showing the pattern of XBP1 direct target gene expression. The top 38 genes that follow a distinct pattern have been displayed. C. Transcriptional regulatory network: Transcription factors that are direct target of XBP1. The genes in the network are differentially expressed (upregulated - red; downregulated - blue) up on 4μ8c treatment. The transcription factors that are not differentially expressed but have a XBP1 ChIPseq peak are shown in the right hand side list.

Despite the preponderance of XBP1’s role in controlling this pathway, other transcription factors are also found to be involved. To examine the regulatory cascade that follows XBP1 regulation, we built a transcriptional regulatory network by extracting annotated transcription factors with promoter or exonic/intronic ChIP-seq peaks (Figure 5C). This network was further complemented by adding differentially expressed genes that have annotated interactions with the target transcription factors in the STRING database (Szklarczyk et al., 2015) (Supplementary Figure S4B).

The transcription factors that are directly regulated by XBP1 can be categorized into three broad functional categories involved in: resolution of protein secretory ER stress, regulation of cell cycle and proliferation, and controlling effector immune cell function. The ER stress involved transcription factors likely facilitate cytokine secretion in Th2 lymphocyte. This prediction is based on the previous reports from secretory cells such as pancreatic acinar cells and plasma cells. These transcription factors, namely Bhlha15, Creb3, Atf6, Atf4, and Creb3l2, have been shown to be involved in secretory stress adaptation of the ER (Acosta-Alvear et al., 2007; Hess et al., 2016; Hetz, 2012; Liang et al., 2006).

The purpose of cell proliferation and cell cycle related transcription factors could be to facilitate the controlled rapid expansion of activated Th2 cells. The immune response related factors are likely involved in Th2 differentiation and cytokine production. Therefore we wanted to test the effect of XBP1s down regulation in cytokine secretion, cell proliferation and cytokine production.

### The IRE1a-XBP1 pathway controls cytokine secretion in T helper cells

The genome-wide comparison of XBP1s regulated genes predicts that the factor is involved in secretion of cytokines. To validate this prediction we blocked IRE1a endonuclease activity in Th2 cells and analyzed the cell culture supernatant to quantify the IL4 level by ELISA. We selected IL4 as a testable candidate cytokine because its mRNA and protein are unchanged by down regulation of XBP1 (Supplementary Figure S5 left panel, Figure 6 left and middle panel of top row). We found that the secretion of IL4 is significantly inhibited in 4μ8c treated cells (Figure 6, right panel of top row). As expected this result supports the involvement of the IRE1a-XBP1 pathway in facilitating cytokine secretion in Th2 cells as predicted. A similar result was observed for IFNγ secretion in Th1 cells (data not shown).

**Figure 6:**
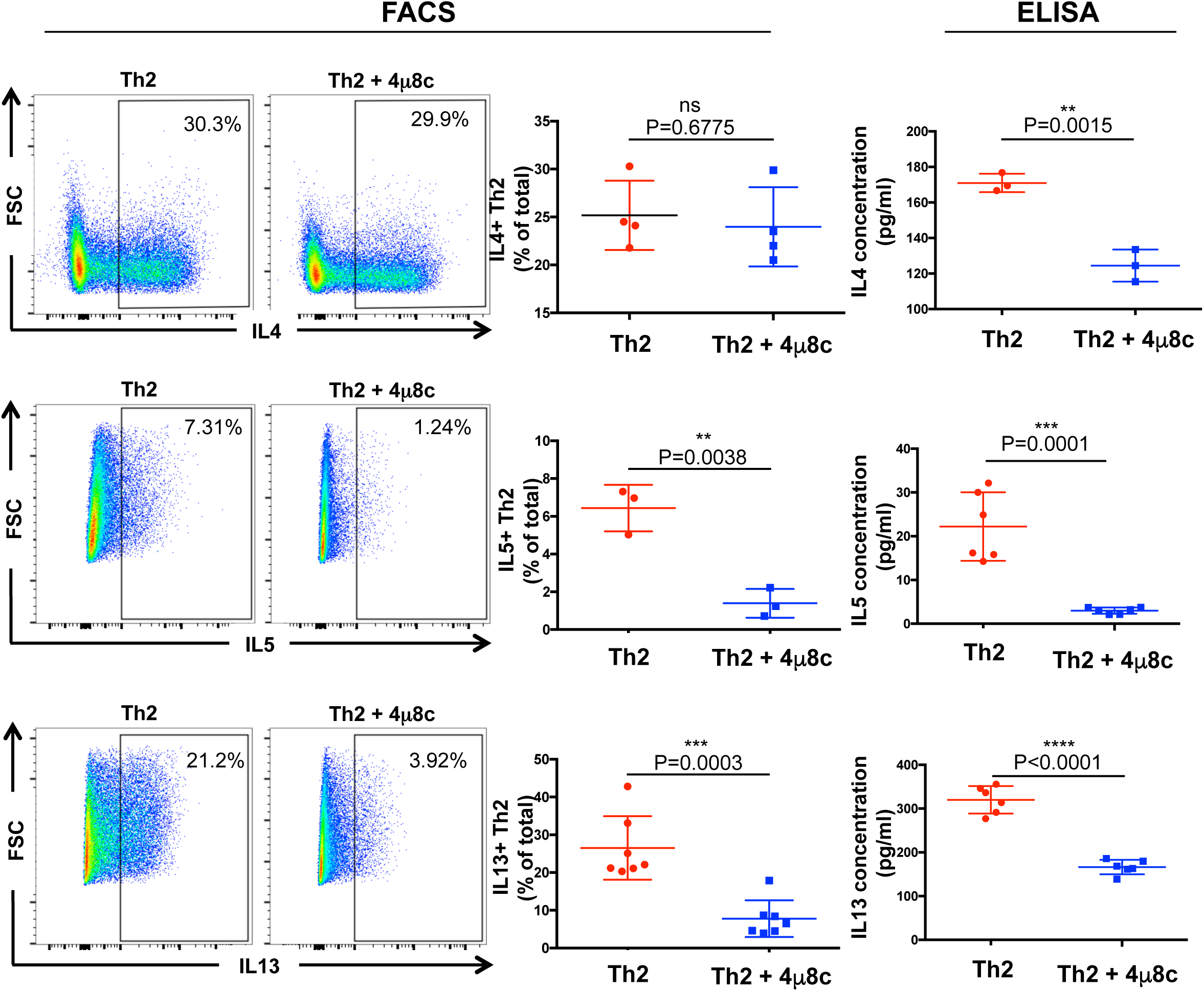
*IRE1a-XBP1 pathway is required for cytokine expression and secretion in Th2 lymphocyte.* Naïve T helper cells were cultured following Th2 activation condition in the presence of IRE1a inhibitor 4μ8c for 3 days, rested for two days, reactivated by coated plate, and analyzed by flow cytometry to detect intra-cellular cytokines IL4, IL5 and IL13 expression. Representative FACS profiles are displayed in the first two columns. The intra-cellular cytokine expression is compared in column 3, with three to seven independent biological replicates. Fourth column: Cell culture supernatants from 4μ8c treated or DMSO treated Th2 were analyzed by ELISA to measure the cytokine concentration.

### IRE1a-XBP1 controls IL13 and IL5 cytokine expression

IL5 and IL13 are two prominent type-2 cytokines that are involved in eosinophilia, allergies and helminth infection. We found that inhibition of IRE1a-XBP1 pathway significantly suppresses the IL5 and IL13 protein expression and secretion into the culture medium (Figure 6 right panels of middle and bottom row). Bioinformatics analysis of the Th2 transcriptome predicts that the IRE1a-XBP1 pathway positively controls IL5 and IL13 gene expression, because both the genes were identified as differentially expressed genes upon IRE1a inhibition (Supplementary Table S1). We validated this prediction by RT-qPCR mediated gene expression analysis (Supplementary Figure 5, middle and right panel) and flow cytometry (Figure 6). These results suggest a transcriptional involvement of the pathway regulating IL5 and IL13. IL5 and IL13 genes are not direct targets of XBP1 transcription factor in the sense of having promoter/genic XBP1 peaks. Hence they must be regulated either from a distal XBP1 binding site or indirectly. Notably the IL4 mRNA and protein levels are not affected indicating specific regulation of IL5 and IL13.

### IRE1a-XBP1 pathway facilitates activation-dependent T helper cell proliferation

Cell proliferation rate is a resultant outcome of the interaction of positive and negative regulators’ interaction. We observed that genes encoding both positive and negative regulators of cell proliferation genes are differentially expressed when the IRE1a-XBP1 pathway was blocked by 4μ8c (Figure 7A, left panel), of which many genes were found to be direct targets of XBP1 (Figure 7A, right panel). This observation predicts a change in proliferation rate upon IRE1a inhibition. Therefore we were interested in checking the effect of IRE1a-XBP1 inhibition on cell proliferation. We performed cell proliferation assay using Th2 cells. We found that down regulation of XBP1s inhibits cell proliferation significantly (Figure 7B), but does not induce cell death (Supplementary Figure S6).

**Figure 7:**
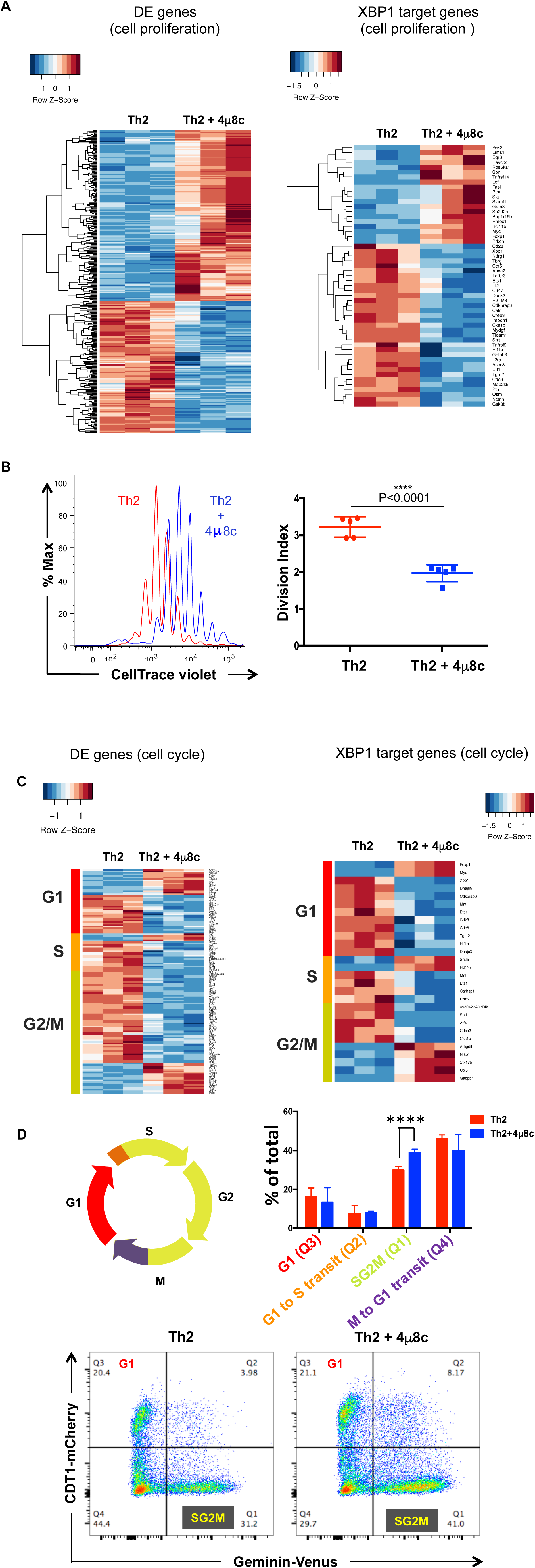
*IRE1a-XBP1 pathway promotes activation dependent Th2 cell proliferation and cell cycling.* A. Left panel: Hierarchical clustering of differentially expressed cell proliferation associated genes in the 4μ8c treated and untreated Th2 transcriptome. Right panel: Hierarchical clustering of XBP1 direct target genes that are known to be involved in cell proliferation. B. Splenic naïve T helper cells were stained with CellTrace Violet dye and activated for 72 hours under Th2 differentiation conditions and analyzed by flow cytometry. Generations of Th2 cells are in “red” and 4μ8c treated cells are in “blue” in the histogram of cell proliferation (left panel, one representative experiment). Graphical representation of division index as obtained from five independent biological replicates (right panel). C. Left panel: Heatmap of differentially expressed cell cycle stage associated genes in the 4μ8c treated and untreated Th2 transcriptome. Right panel: Heatmap of XBP1 direct target genes that are known to be involved in cell cycling. D. Cell cycle analysis of Th2 lymphocytes after 72 hours of activation, using FUCCI mouse line that express mCherry tagged CDT1 and Venus tagged Geminin. Upper left: Diagrammatic representation of cell cycle stages in used FUCCI mouse. Upper right: Comparison of cells (% of total) obtained from different stages of cell cycle in Th2 and 4μ8c treated Th2 (n=6). Lower panels: One representative FACS profile of Th2 and 4μ8c treated Th2 showing CDT1 and Geminin expressing cells.

T helper cell proliferation is associated with differentiation and cytokine production. The reduced IL5 and IL13 expression (Figure 6) could potentially be explained by the fact that cell proliferation is retarded. However, if reduced proliferation was the primary reason for lack of secretion, IL4 production would also be inhibited. Yet, we observed no significant change in IL4 expression upon IRE1a inhibition (Figure 6, Supplementary Figure S5). To examine this discrepancy further, we performed cell proliferation assays using IL13-GFP and IL4-GFP reporter mouse lines. In IL4-GFP expressing Th2 cells we observed an inhibition of IL4 production in the first few generations of cell division up to 72 hours upon 4μ8c treatment (Supplementary Figure S7). But after five cell divisions the difference in IL4 expression becomes insignificant. This observation suggests that the retardation of proliferation due to the IRE1a inhibition is not sufficient to inhibit IL4 expression. In contrast, in IL13-GFP we observed the decrease in IL13 expression from the very first generation and this continues throughout the later generations (Supplementary Figure S8).

### IRE1a inhibition delays cell cycle progression through the S and G2/M phase

Bioinformatics analysis of differentially expressed genes (Th2 vs 4μ8c treated Th2) and XBP1 direct target genes reveals several genes that are involved in controlling cell cycle progression through different stages (i.e. G1, S, G2/M) were clustered in to two groups up or down regulated (Figure 7C). We took genes differentially expressed in 4m8c treated Th2 compared to untreated Th2 (adjusted p-value <0.05) (Figure 7C, left) and the genes differentially expressed XBP1 direct target genes (Figure 7C, right), and checked for known roles across distinct cell cycle stages using either a manually curated list based on RNA-seq data or published database (Santos et al., 2015). We found many genes from all cell cycle stages (i.e. G1, S and G2/M) were affected. To identify the cell cycle stage that is regulated by IRE1a-XBP1 pathway we exploited a genetically modified mouse line (i.e. <u>F</u>luorescence <u>U</u>biquitin <u>C</u>ell *C*ycle <u>I</u>ndicator (FUCCI) mouse) that expresses mCherry tagged CDT1 protein and Venus tagged Geminin protein. The mouse line faithfully reports on the cell cycle state with mCherry expressed in G1, Venus expressed in S and G2M phase, and two transient phases without any fluorescence during M to G1 and G1 to S transition (Figure 7D, upper left panel). We compared cell cycle profiles of vehicle and 4μ8c treated Th2 cells during T cell activation. We found that cells accumulated in the S and/or G2/M phase when the IRE1a-XBP1 pathway is blocked (Figure 7D). Similar results were obtained in a different approach using BrdU incorporation assay with DAPI staining (Supplementary Figure S9).

## Conclusions

The primary aim of this study was to investigate the role of XBP1 transcription factor in Th2 lymphocytes; to identify the Th2 specific XBP1 target genes and their involvement regulating Th2 cell biology. We showed evidence that the IRE1a-XBP1 pathway is engaged in resolving secretory stress to meet robust cytokine synthesis and secretion, and controls multiple important cellular properties of T helper lymphocyte. It regulates activation dependent T helper cell proliferation and cytokine production, the two key features of T helper cell during activation. The study revealed a large transcriptional regulatory network governed by XBP1. The comprehensive repertoire of XBP1 regulated genes and its genome-wide binding map provides a valuable resource for future work. We built a transcriptional regulatory map by integrating XBP1 ChIPmentation and RNAseq data, which portrays the bigger picture of the involvement of the XBP1 transcription factor in regulating target genes including other transcription factors. To visualize the data we created an easily browsable online database: http://data.teichlab.org

ER-stress is known to be involved in several pathological situations. The pathway promotes cancer progression by providing metabolic advantage to the neoplastic cancer cells to acclimatize to the stressed tumor microenvironment. During the antitumor immune response, the XBP1 pathway induces tolerance in DCs. The pathway promotes asthmatic, allergic and eosinophilic immune reactions and is involved in immunometabolism of macrophages in obesity. The pathway can be modulated by drug such as 4μ8c and STF-083010 and is under intensive investigation. Further studies will have to be carried out to determine whether the modulation of the pathway can bring patients benefit. This study shows evidence that perturbation of the IRE1a-XBP1 pathway may interfere with normal physiological activation of Th2 and could be exploited in settings where Th2 lymphocytes are pathologic such as asthma, allergies and eosinophilia. Two prominent cytokines, IL5 and IL13, which promote allergies and eosinophilia, are under the control of IRE1a-XBP1 pathway in Th2 lymphocytes. In future, locus specific mechanistic dissection of the XBP1 mediated transcription process in Th2 lymphocytes, and *in vivo* immunobiological studies on novel Th2 specific XBP1 target genes are required to understand how the XBP1 transcription factor orchestrates locus control and to what extent it controls Th2 mediated immune responses.

## Materials and Methods

### Materials

CD4^+^CD62L^+^ T Cell Isolation Kit II, mouse (Miltenyi Biotec, 130-093-227)

Naive CD4^+^ T Cell Isolation Kit, mouse (Miltenyi Biotec, 130-104-453)

FITC BrdU Flow Kit (BD Pharmingen, 51-2354AK)

Mouse IL-13 ELISA Ready-SET-Go Kit (eBioscience, 88-7137-22)

Mouse IL-4 ELISA Ready-SET-Go Kit (eBioscience, 88-7044-88)

Mouse IL-5 ELISA (BD Biosciences, 555236)

PE Mouse anti-XBP1S Clone Q3-695 (BD Pharmingen, 562642)

XBP1 (M-186)X-(Santa cruz, Sc 7160x)

IL5-PE (BD Pharmingen, 554395)

IL4-APC, Clone 11B11 (eBioscience, 17-7041-82)

IL13-AF488, Clone eBio3A (eBioscience, 53-7133-82)

IFNg-Per CP Cy5.5, Clone XMG1.2 (eBioscience, 45-7311-82)

FACS Staining buffer (eBioscience, 00-4222-26)

IC Fixation buffer (eBioscience, 00-8222-49)

Fixation/Permeabilization diluent (eBioscience, 00-5223-56)

Fixation/Permeabilization concentrate (eBioscience, 00-5123-43)

Permeabilization buffer (eBioscience, 00-8333-56)

SV Total RNA Isolation System (Promega, Z3101)

Transcriptor High Fidelity cDNA Synthesis kit (Roche, 05081955001)

SYBR™ Select Master Mix (Applied Biosystems, 4472908).

Western blot Antibodies

IRE1 α (14C10) Rabbit mAb (Cell Signaling, #3294)

IRE1alpha [p Ser724] Antibody (Novus biologicals,NB100-2323)

### Mice

Animal research at WTSI and MRC-LMB was conducted under license from the UK Home Office (PPLH 70/7968 and 70/8381, respectively) and the institute’s animal welfare and ethical review body approved used protocols. The mice (C57BL/6, IL13-eGFP reporter, IL4-eGFP reporter and FUCCI) were maintained under specific pathogen-free conditions at the Wellcome Trust Genome Campus Research Support Facility (Cambridge, UK) and were used at 6-12 weeks of age. All procedures were in accordance with the Animals (Scientific Procedures) Act 1986.

### T helper Cell Culture

Splenic naive T helper cells were purified with the CD4^+^CD62L^+^T Cell Isolation Kit II (Miltenyi Biotec) and polarized *in vitro* toward differentiated Th2 subtype as described before in (Mahata et al., 2014). In brief, naive cells were seeded into anti-CD3e (2 μg/ml, clone 145-2C11, eBioscience) and anti-CD28 (5 μg/ml, clone 37.51, eBioscience) antibody coated 96-well round bottom plates. The medium contained the following cytokines and/or antibodies for Th2 subtype, recombinant murine IL-2 (10 ng/ml, R&D Systems), recombinant murine IL-4 (10 ng/ml,
R&D Systems) and neutralizing anti-IFN-g (5μg/ml, cloneXMG1.2eBioscience). The cells were removed from the activation plate on day 4 (72 hours). Th2 cells were cultured for another two days in the absence of CD3 and CD28 stimulation. Then, cells were restimulated by coated plate for 6 hrs. For flow cytometric detection cells were treated with monensin (2μM, eBioscience) for the last 3 hours.

### Reverse Transcription Quantitative PCR (RT-qPCR)

Total RNA was isolated from 2 millions cells by SV total RNA isolation kit (Promega). cDNA was prepared by annealing 500ng RNA with oligo dT as per manufacturers instructions (Transcriptor High Fidelity cDNA Synthesis kit, Roche).The cDNA samples were diluted 10 times with H20. 2 μl of cDNA was used in 12 μl qPCR reactions with appropriate primers and SYBR Green PCR Master Mix (Applied Biosystems). Experiments were performed at least 3 times and data represent mean values +/− standard deviation. For XBP1 mRNA was amplified by PCR and products were separated by electrophoresis through a 2.5% agarose gel and visualized by ethidium bromide staining. The primer list is provided below:

IL4-F: 5’ - AACTCCATGCTTGAAGAAGAACTC-3’

IL4-R: 5’ - CCAGGAAGTCTTTCAGTGATGTG -3’

IL13-F: 5’ - CCTGGCTCTTGCTTGCCTT-3’

IL13-R: 5’ - GGTCTTGTGTGATGTTGCTCA-3’

IL5-F: 5’ - GCAATGAGACGATGAGGCTTC-3’

IL5-R: 5’ - CCCCTGAAAGATTTCTCCAATG-3’

ERN1-F: 5’ - ACACCGACCACCGTATCTCA-3’

ERN1-R: 5’ - CTCAGGATAATGGTAGCCATGTC-3’

XBP1 -F: 5’ - ACACGCTTGGGAATGGACAC-3’

XBP1-R: 5’- CCATGGGAAGATGTTCTGGG-3’

RPLP0-F: 5’- CACTGGTCTAGGACCCGAGAA-3’

RPLP0-R: 5’- GGTGCCTCTGGAGATTTTCG-3’

### ELISA

IL13, IL4 and IL5 concentration in the Th2 culture supernatants were quantified using ELISA kit following manufacturer’s instruction (see “Materials” for the kit specification).

### Flow cytometry

In worm infection mouse experiments, splenocytes were prepared on day 7 postinfection from N. brasiliensis infected or control uninfected mice, stained with anti-CD3e, anti-CD4 (eBioscience), and XBP1s-PE (BD Pharmingen) antibodies following the mouse regulatory T Cell staining kit protocol (eBioscience), and were measured by flow cytometry on a Fortessa (BD Biosciences) using FACSDiva. The data were analyzed by the FlowJo software. For *in vitro* Th cell experiments, staining was performed following eBioscience intracellular staining protocol for cytokines and nuclear staining/transcription factor staining protocol for XBP1 transcription factor using eBioscience reagents and kit protocol. The following antibodies were fluorescent dye-conjugated primary antibodies: IL-4, IL-13, IL-5, CD4 and IFNγ (eBioscience), XBP1s (BD Pharmingen). Stained cells were analyzed on a Fortessa (BD Biosciences) using FACSDiva and FlowJo software. CompBeads (BD Biosciences) were used for compensation where distinct positively stained populations were unavailable.

### Cell Proliferation Assay

Naive Th cells were stained with CellTrace Violet following the CellTrace Violet Cell Proliferation Kit (Invitrogen) protocol and cultured under activation-differentiation conditions for Th2 as described previously, in the presence or absence of 15μM 4μ8c for 4 days. Flow cytometry was performed using a BD Fortessa and data analysis with FlowJo software.

### N. brasiliensis infection and splenocyte preparation

C57BL/6 female mice were subcutaneously injected with 100 μl (300/500 live third stage *N. brasiliensis* larvae per dose). Spleen was taken from infected mice 7 days after infection. Cells were isolated from spleen by smashing the tissue though a 70μιτι cell strainer and suspended in RBC lysis buffer (eBioscience). Single cell suspensions of splenocytes were then stained following FACS staining protocol.

### Analysis of bulk RNA-sequencing data

For each sample, reads were mapped to the *Mus musculus* genome (GRCm38) using GSNAP with default parameters (Wu and Nacu, 2010). Uniquely mapped reads to the genome were counted using htseq-count (http://www-huber.embl.de/users/anders/HTSeq/) and normalized with size factors calculated by DESeq2 (Anders and Huber, 2010). Differentially expressed genes across conditions were identified using DESeq2 with an adjusted p-value cutoff < 0.05.

### In vivo *XBP1 expression analysis in single cells*

Reference sequences (Dec. 2011 (GRCm38/mm10)) were downloaded from UCSC refSeq at: http://hgdownload.soe.ucsc.edu/goldenPath/mm10/bigZips/refMrna.fa.gz Raw fastq files are downloaded from ENA (www.ebi.ac.uk/ena). Cell annotations were downloaded from ArrayExpress (www.ebi.ac.uk/arrayexpress). The refSeq Ids for the spliced and unspliced forms of XBP1 are NM_001271730 and NM_013842 respectively. The reads were mapped with salmon version 0.8.2 by kmer=21, and transcript per million (TPM) were used for quantification. Accession codes: E-MTAB-4619 (Proserpio et al., 2016).

### XBP1 ChIPmentation

20 million cells from each sample were crosslinked in 1% HCHO (prepared in 1X DPBS) at room temperature for 10 minutes, and HCHO was quenched by the addition of glycine at a final concentration of 0.125 M. Cells were pelleted at 4°C at 2000 × g, washed with ice-cold 1X DPBS twice, and snapped frozen in liquid nitrogen. The cell pellets were stored in −80°C until the experiments were performed. ChIPmentation was performed according to the version 1.0 of the published protocol (Schmidl et al., 2015b) with some modifications at the ChIP stage. Briefly, cell pellets were thawed on ice, and lysed in 300 μl ChIP Lysis Buffer I (50 mM HEPES.KOH, pH 7.5, 140 mM NaCl, 1 mM EDTA, pH 8.0, 10% Glycerol, 0.5% NP-40, 0.25% Triton X-100) on ice for 10 minutes. Then cells were pelleted at 4°C at 2000 x g for 5 minutes, and washed by 300 μl ChIP Lysis Buffer II (10 mM Tris.Cl, pH 8.0, 200 mM NaCl, 1 mM EDTA, pH 8.0, 0.5 mM EGTA, pH 8.0), and pelleted again at 4°C at 2000 × g for 5 minutes. Nuclei were resuspended in 300 μl ChIP Lysis Buffer III (10 mM Tris.Cl, pH 8.0, 100 mM NaCl, 1 mM EDTA, 0.5 mM EGTA, 0.1% Sodium Deoxycholate, 0.5% N-Lauryolsarcosine). Chromatin was sonicated using Bioruptor Pico (Diagenode) with 30 seconds ON/30 seconds OFF for 5 cycles. 30 μl 10% Triton X-100 were added into each sonicated chromatin, and insoluble chromatin was pelleted at 16,100 x g at 4°C for 10 minutes. 1 μl supernatant was taken as input control. The rest of the supernatant was incubated with 10 μl Protein A Dynabeads (Invitrogen) pre-bound with 1 μg XBP1 antibody (XBP1 (M-186)X - Santa cruz), in a rotating platform in a cold room overnight. Each immunoprecipitation (IP) was washed with 500 μl RIPA Buffer (50 mM HEPES.KOH, pH 7.5, 500 mM LiCl, 1 mM EDTA, 1% NP-40, 0.7% Sodium Deoxycholate, check components) for 3 times. Then, each IP was washed with 500 μl 10 mM Tris, pH 8.0 twice, and resuspended in 30 μl tagmentation reaction mix (10 mM Tris.Cl, pH 8.0, 5 mM Mg2Cl, 1 μl TDE1 (Nextera)). Then, the tagmentation reaction was put on a thermomixer at 37°C for 10 minutes at 800 rpm shaking. After the tagmentation reaction, each IP was washed sequentially with 500 μl RIPA Buffer twice, and 1X TE NaCl (10 mM Tris.Cl, pH 8.0, 1 mM EDTA, pH 8.0, 50 mM NaCl) once. Elution and reverse-crosslinking was done by resuspending the beads with 100 μl ChIP Elution Buffer (50 mM Tris.Cl, pH 8.0, 10 mM EDTA, pH 8.0, 1% SDS) on a thermomixer at 65°C overnight, 1,400 rpm. DNA was purified by MinElute PCR Purification Kit (QIAGEN, cat no. 28004 and eluted in 12.5 μl Buffer EB (QIAGEN kit, cat no 28004), which yielded ~10 μl ChIPed DNA.

#### The library preparation reactions contained the following

10 μl purified DNA (from above), 2.5 μl PCR Primer Cocktails (Nextera DNA Library Preparation Kit, Illumina Cat no. FC-121-1030), 2.5 μl N5xx (Nextera Index Kit, Illumina cat no. FC-121-1012), 2.5 μl N7xx (Nextera index kit, Illumina cat no. FC-121-1012), 7.5 μl NPM PCR Master Mix (Nextera DNA Library Preparation Kit, Illumina Cat no. FC-121-1030)

#### PCR was set up as follows

72°C, 5 mins; 98°C, 2 mins; [98°C, 10 secs, 63°C, 30 secs, 72°C, 20 secs] x 12; 10°C hold The amplified libraries were purified by double AmpureXP beads purification: first with 0.5X bead ratio, keep supernatant, second with 1.4X bead ratio, keep bound DNA. Elution was done in 20 μl Buffer EB (QIAGEN).

1 μl of library was run on an Agilent Bioanalyzer to see the size distribution. Sequencing was done on an Illumina Hiseq2000 platform using the v4 chemistry (75 bp PE).

### ChIPmentation analysis

The reads were first trimmed using Trimmomatic 0.3664 with settings ILLUMINACLIP:NexteraPE-PE.fa:2:30:10 LEADING:3 TRAILING:3

SLIDINGWINDOW:4:15 MINLEN:30. Peaks were then called using MACS265, merged over time, and annotated using HOMER66.

The quality of the peaks was assessed using the two available replicates of XBP1. *Inferred regulatory cascade of XBP1*:

Transcription factors were obtained from the AnimalTFDB 2.0 (Zhang et al., 2015), and were defined as targets of XBP1 if they were intersected by a ChIPmentation peak and differentially expressed between Th2 (control) and 4μ8c treated Th2. Genes were defined as targeted by these transcription factors (Supplementary Figure S4) if in the STRING version 10 database (Szklarczyk et al., 2015) they had an “expression” mode of interaction with a score greater than 200 with these transcription factors in mouse, and were differentially expressed between Th2 (control) and 4μ8c treated Th2.

## Acknowledgements

We would like to thank Kerstin Meyer, for useful discussions and comments on the manuscript, Helen E. Jolin for technical help with *in vivo* experiments; Padraic Fallon for providing *N. brasiliensis* larvae; Bee Ling Ng, Chris Hall and Jennie Graham for help with flow cytometry; Research Support Facility, WTSI, for their technical help and animal husbandry.

## References

Acosta-Alvear, D., Zhou, Y., Blais, A., Tsikitis, M., Lents, N.H., Arias, C., Lennon, C.J., Kluger, Y., and Dynlacht, B.D. (2007). XBP1 controls diverse cell type- and condition-specific transcriptional regulatory networks. Molecular cell 27, 53-66.

Anders, S., and Huber, W. (2010). Differential expression analysis for sequence count data. Genome biology 11, R106.

Bettigole, S.E., and Glimcher, L.H. (2015). Endoplasmic reticulum stress in immunity. Annual review of immunology 33, 107-138.

Bettigole, S.E., Lis, R., Adoro, S., Lee, A.H., Spencer, L.A., Weller, P.F., and Glimcher, L.H. (2015). The transcription factor XBP1 is selectively required for eosinophil differentiation. Nature immunology 16, 829-837.

Bird, J.J., Brown, D.R., Mullen, A.C., Moskowitz, N.H., Mahowald, M.A., Sider, J.R., Gajewski, T.F., Wang, C.R., and Reiner, S.L. (1998). Helper T cell differentiation is controlled by the cell cycle. Immunity 9, 229-237.

Brucklacher-Waldert, V., Ferreira, C., Stebegg, M., Fesneau, O., Innocentin, S., Marie, J.C., and Veldhoen, M. (2017). Cellular Stress in the Context of an Inflammatory Environment Supports TGF-beta-Independent T Helper-17 Differentiation. Cell reports 19, 2357-2370.

Brunsing, R., Omori, S.A., Weber, F., Bicknell, A., Friend, L., Rickert, R., and Niwa, M. (2008). B- and T-cell development both involve activity of the unfolded protein response pathway. The Journal of biological chemistry 283, 17954-17961.

Calfon, M., Zeng, H., Urano, F., Till, J.H., Hubbard, S.R., Harding, H.P., Clark, S.G., and Ron, D. (2002). IRE1 couples endoplasmic reticulum load to secretory capacity by processing the XBP-1 mRNA. Nature 415, 92-96.

Camberis, M., Le Gros, G., and Urban, J., Jr. (2003). Animal model of Nippostrongylus brasiliensis and Heligmosomoides polygyrus. Current protocols in immunology / edited by John E Coligan [et al] *Chapter 19,* Unit 19 12.

Chen, X., Iliopoulos, D., Zhang, Q., Tang, Q., Greenblatt, M.B., Hatziapostolou, M., Lim, E., Tam, W.L., Ni, M., Chen, Y., et al. (2014). XBP1 promotes triple-negative breast cancer by controlling the HIF1alpha pathway. Nature 508, 103-107.

Chen, X., Muller, G.A., Quaas, M., Fischer, M., Han, N., Stutchbury, B., Sharrocks, A.D., and Engeland, K. (2013). The forkhead transcription factor FOXM1 controls cell cycle-dependent gene expression through an atypical chromatin binding mechanism. Molecular and cellular biology 33, 227-236.

Cross, B.C., Bond, P.J., Sadowski, P.G., Jha, B.K., Zak, J., Goodman, J.M., Silverman, R.H., Neubert, T.A., Baxendale, I.R., Ron, D., et *al.* (2012). The molecular basis for selective inhibition of unconventional mRNA splicing by an IRE1-binding small molecule. Proceedings of the National Academy of Sciences of the United States of America 109, E869-878.

Cubillos-Ruiz, J.R., Bettigole, S.E., and Glimcher, L.H. (2017). Tumorigenic and Immunosuppressive Effects of Endoplasmic Reticulum Stress in Cancer. Cell 168, 692-706.

Cubillos-Ruiz, J.R., Silberman, P.C., Rutkowski, M.R., Chopra, S., Perales-Puchalt, A., Song, M., Zhang, S., Bettigole, S.E., Gupta, D., Holcomb, K., et al. (2015). ER Stress Sensor XBP1 Controls Anti-tumor Immunity by Disrupting Dendritic Cell Homeostasis. Cell 161, 1527-1538.

Ellyard, J.I., Simson, L., and Parish, C.R. (2007). Th2-mediated anti-tumour immunity: friend or foe? Tissue antigens 70, 1-11.

Frakes, A.E., and Dillin, A. (2017). The UPRER: Sensor and Coordinator of Organismal Homeostasis. Molecular cell 66, 761-771.

Gett, A.V., and Hodgkin, P.D. (1998). Cell division regulates the T cell cytokine repertoire, revealing a mechanism underlying immune class regulation. Proceedings of the National Academy of Sciences of the United States of America 95, 9488-9493.

Grootjans, J., Kaser, A., Kaufman, R.J., and Blumberg, R.S. (2016). The unfolded protein response in immunity and inflammation. Nature reviews Immunology 16, 469-484.

Heinz, S., Benner, C., Spann, N., Bertolino, E., Lin, Y.C., Laslo, P., Cheng, J.X., Murre, C., Singh, H., and Glass, C.K. (2010). Simple combinations of lineage-determining transcription factors prime cis-regulatory elements required for macrophage and B cell identities. Molecular cell 38, 576-589.

Hess, D.A., Strelau, K.M., Karki, A., Jiang, M., Azevedo-Pouly, A.C., Lee, A.H., Deering, T.G., Hoang, C.Q., MacDonald, R.J., and Konieczny, S.F. (2016). MIST1 Links Secretion and Stress as Both Target and Regulator of the UPR. Molecular and cellular biology.

Hetz, C. (2012). The unfolded protein response: controlling cell fate decisions under ER stress and beyond. Nature reviews Molecular cell biology 13, 89-102.

Hetz, C., Chevet, E., and Harding, H.P. (2013). Targeting the unfolded protein response in disease. Nature reviews Drug discovery 12, 703-719.

Hetz, C., Martinon, F., Rodriguez, D., and Glimcher, L.H. (2011). The unfolded protein response: integrating stress signals through the stress sensor IRE1alpha. Physiological reviews 91, 1219-1243.

Hetz, C., and Papa, F.R. (2017). The Unfolded Protein Response and Cell Fate Control. Molecular cell.

Hosokawa, H., Tanaka, T., Kato, M., Shinoda, K., Tohyama, H., Hanazawa, A., Tamaki, Y., Hirahara, K., Yagi, R., Sakikawa, I., et al. (2013). Gata3/Ruvbl2 complex regulates T helper 2 cell proliferation via repression of Cdkn2c expression. Proceedings of the National Academy of Sciences of the United States of America 110, 18626-18631.

Hotamisligil, G.S. (2010). Endoplasmic reticulum stress and the inflammatory basis of metabolic disease. Cell 140, 900-917.

Iwakoshi, N.N., Pypaert, M., and Glimcher, L.H. (2007). The transcription factor XBP-1 is essential for the development and survival of dendritic cells. The Journal of experimental medicine 204, 2267-2275.

Janssens, S., Pulendran, B., and Lambrecht, B.N. (2014). Emerging functions of the unfolded protein response in immunity. Nature immunology 15, 910-919.

Kaser, A., and Blumberg, R.S. (2010). Survive an innate immune response through XBP1. Cell research 20, 506-507.

Lee, A.H., Iwakoshi, N.N., and Glimcher, L.H. (2003). XBP-1 regulates a subset of endoplasmic reticulum resident chaperone genes in the unfolded protein response. Molecular and cellular biology 23, 7448-7459.

Liang, G., Audas, T.E., Li, Y., Cockram, G.P., Dean, J.D., Martyn, A.C., Kokame, K., and Lu, R. (2006). Luman/CREB3 induces transcription of the endoplasmic reticulum (ER) stress response protein Herp through an ER stress response element. Molecular and cellular biology 26, 7999-8010.

Mahata, B., Zhang, X., Kolodziejczyk, A.A., Proserpio, V., Haim-Vilmovsky, L., Taylor, A.E., Hebenstreit, D., Dingler, F.A., Moignard, V., Gottgens, B., et al. (2014). Singlecell RNA sequencing reveals T helper cells synthesizing steroids de novo to contribute to immune homeostasis. Cell reports 7, 1130-1142.

Mathelier, A., Fornes, O., Arenillas, D.J., Chen, C.Y., Denay, G., Lee, J., Shi, W., Shyr, C., Tan, G., Worsley-Hunt, R., et al. (2016). JASPAR 2016: a major expansion and update of the open-access database of transcription factor binding profiles. Nucleic acids research 44, D110-115.

McLean, C.Y., Bristor, D., Hiller, M., Clarke, S.L., Schaar, B.T., Lowe, C.B., Wenger, A.M., and Bejerano, G. (2010). GREAT improves functional interpretation of cis-regulatory regions. Nature biotechnology 28, 495-501.

Muller, G.A., Quaas, M., Schumann, M., Krause, E., Padi, M., Fischer, M., Litovchick, L., DeCaprio, J.A., and Engeland, K. (2012). The CHR promoter element controls cell cycle-dependent gene transcription and binds the DREAM and MMB complexes. Nucleic acids research 40, 1561-1578.

Murphy, K.M., and Reiner, S.L. (2002). The lineage decisions of helper T cells. Nature reviews Immunology 2, 933-944.

Neill, D.R., Wong, S.H., Bellosi, A., Flynn, R.J., Daly, M., Langford, T.K., Bucks, C., Kane, C.M., Fallon, P.G., Pannell, R., et al. (2010). Nuocytes represent a new innate effector leukocyte that mediates type-2 immunity. Nature 464, 1367-1370.

Osorio, F., Tavernier, S.J., Hoffmann, E., Saeys, Y., Martens, L., Vetters, J., Delrue, I., De Rycke, R., Parthoens, E., Pouliot, P., et al. (2014). The unfolded-protein-response sensor IRE-1alpha regulates the function of CD8alpha+ dendritic cells. Nature immunology 15, 248-257.

Proserpio, V., Piccolo, A., Haim-Vilmovsky, L., Kar, G., Lonnberg, T., Svensson, V., Pramanik, J., Natarajan, K.N., Zhai, W., Zhang, X., et al. (2016). Single-cell analysis of CD4+ T-cell differentiation reveals three major cell states and progressive acceleration of proliferation. Genome biology 17, 103.

Santos, A., Wernersson, R., and Jensen, L.J. (2015). Cyclebase 3.0: a multi-organism database on cell-cycle regulation and phenotypes. Nucleic acids research 43, D1140-1144.

Schmidl, C., Rendeiro, A.F., Sheffield, N.C., and Bock, C. (2015a). ChIPmentation: fast, robust, low-input ChIP-seq for histones and transcription factors. Nature methods 12, 963-965.

Schmidl, C., Rendeiro, A.F., Sheffield, N.C., and Bock, C. (2015b). ChIPmentation: fast, robust, low-input ChIP-seq for histones and transcription factors. Nature methods 12, 963-965.

Shan, B., Wang, X., Wu, Y., Xu, C., Xia, Z., Dai, J., Shao, M., Zhao, F., He, S., Yang, L., et al. (2017). The metabolic ER stress sensor IRE1alpha suppresses alternative activation of macrophages and impairs energy expenditure in obesity. Nature immunology 18, 519-529.

Szklarczyk, D., Franceschini, A., Wyder, S., Forslund, K., Heller, D., Huerta-Cepas, J., Simonovic, M., Roth, A., Santos, A., Tsafou, K.P., et al. (2015). STRING v10: proteinprotein interaction networks, integrated over the tree of life. Nucleic acids research 43, D447-452.

Thaxton, J.E., Wallace, C., Riesenberg, B., Zhang, Y., Paulos, C.M., Beeson, C.C., Liu, B., and Li, Z. (2017). Modulation of Endoplasmic Reticulum Stress Controls CD4+ T-cell Activation and Antitumor Function. Cancer immunology research 5, 666-675.

Todd, D.J., McHeyzer-Williams, L.J., Kowal, C., Lee, A.H., Volpe, B.T., Diamond, B., McHeyzer-Williams, M.G., and Glimcher, L.H. (2009). XBP1 governs late events in plasma cell differentiation and is not required for antigen-specific memory B cell development. The Journal of experimental medicine 206, 2151-2159.

Walker, J.A., and McKenzie, A.N.J. (2017). TH2 cell development and function. Nature reviews Immunology.

Wegmann, T.G., Lin, H., Guilbert, L., and Mosmann, T.R. (1993). Bidirectional cytokine interactions in the maternal-fetal relationship: is successful pregnancy a TH2 phenomenon? Immunology today 14, 353-356.

Wu, T.D., and Nacu, S. (2010). Fast and SNP-tolerant detection of complex variants and splicing in short reads. Bioinformatics 26, 873-881.

Zhang, H.M., Liu, T., Liu, C.J., Song, S., Zhang, X., Liu, W., Jia, H., Xue, Y., and Guo, A.Y. (2015). AnimalTFDB 2.0: a resource for expression, prediction and functional study of animal transcription factors. Nucleic acids research 43, D76-81.

Zhang, Y., Liu, T., Meyer, C.A., Eeckhoute, J., Johnson, D.S., Bernstein, B.E., Nusbaum, C., Myers, R.M., Brown, M., Li, W., et al. (2008). Model-based analysis of ChIP-Seq (MACS). Genome biology 9, R137.

Zhu, J., Yamane, H., and Paul, W.E. (2010). Differentiation of effector CD4 T cell populations (*). Annual review of immunology 28, 445-489.

